# The Alphaherpesvirus Latency Associated Promoter 2 (LAP2) Drives Strong Transgene Expression in Peripheral Tissue Depending on Administration Route and AAV Serotype

**DOI:** 10.1101/2022.08.04.502832

**Authors:** Carola J. Maturana, Angela Chan, Jessica L. Verpeut, Esteban A. Engel

**Affiliations:** Princeton Neuroscience Institute, Princeton University, Princeton, NJ 08544, USA; Department of Psychology, Arizona State University, Tempe, AZ 85287, USA

**Keywords:** AAV8, AAV9, Gene Therapy, Viral vector, Recombinant AAV, Gene Promoter, LAP, Latency-Associated Promoter, LAT, Latency-Associated Transcript, Gene Delivery, Liver, Lung, Kidney, Skeletal Muscle, Retro-Orbital Injection, Intramuscular injection

## Abstract

Adeno-associated virus (AAV) has shown great translational potential in treating a variety of diseases often requiring strong and ubiquitous transgene expression. However, the genetic payload of AAV vectors is limited to <4.9 kb and some commonly used gene promoters are large in sizeable and susceptible to transcriptional silencing. We validated a short (404 bp), strong and persistent promoter obtained from the genome of pseudorabies virus (PRV) called alphaherpesvirus latency-associated promoter 2 (LAP2). We evaluated the biodistribution and potency of transgene expression in mouse peripheral tissue and organs when using AAV8-LAP2 and AAV9-LAP2, both of which achieved transgene expression like that of the ubiquitous promoter, EF1α. LAP2 drives potent transgene expression in liver and kidney after systemic retro-orbital administration and in skeletal muscle after intramuscular delivery. Additionally, we observed broad transduction throughout the lung albeit at lower levels than other tissues. Notably, in skeletal muscle LAP2 resulted in preferential transduction of myofiber types 2. A direct side-by-side comparison between LAP2 and EF1α, demonstrates that regardless of the AAV serotype and route of administration, LAP2 is as powerful and persistent as EF1α promoter despite being 66% smaller in size, thus allowing for larger therapeutic payloads.

## Introduction

Viral vectors have emerged as an effective gene transfer platform for research, gene therapy clinical trials, and vaccine development.^1,2^ Different viral vectors have distinct safety profiles, tropism and packaging capacity. Adeno-associated virus (AAV) is safe, exhibits a favorable immune response compared to other vectors, and is generally non-pathogenic. However, it has a rather limited cargo capacity of <4.9 kb^3^, that is further reduced by half (∼2.4 kb) in self-complementary (double-stranded DNA) AAV vectors.^4^ Approaches to increase AAV payload and broaden the spectrum of therapeutic applications include the use of truncated transgenes AAV co-transduction (dual or triple vector strategy).^5,6^ Additionally, minimal sequence design of constitutive promoters has been widely used,^7^ such as miniature chicken β-actin (CBA; 800 bp) or modified miniature CAG promoter (CBh; 800 bp), however these small promoters are susceptible to transcriptional silencing over time.^8,9^ In an effort to improve AAV vectors as a gene therapy tool, we used a short, potent and stable promoter obtained from the genome of the herpesvirus pseudorabies virus (PRV) called alphaherpesvirus latency-associated promoter (LAP). PRV LAP has been shown to remain active during both lytic and latent phases, chronically driving transcription of latency-associated transcripts (LATs).^10^ PRV LAP contains a cluster of two tandem promoters sequences, PRV LAP1 and PRV LAP2 that drive LAT gene expression *in vitro* and *in vivo*.^11,12^ We previously validated three versions of these promoters called LAP1, LAP2 and tandem LAP1_LAP2 and found that there are persistent non-repressible gene promoters with strong and long-lasting pan-neuronal expression in the brain and spinal cord after AAV transduction.^13^ Previous research administering PRV LAP by intracranial inoculation in mice have shown spread to peripheral organs such as liver and lung, suggesting a putative role of LAP in PRV infectivity.^14^ Furthermore, Cheung and Smith in 1999, suggested that PRV-LAP2 controlled gene expression in both neuronal and non-neuronal cells, in contrast to PRV-LAP1 which was active only in neuronal cells.^11^ Given that LAP2 (404 bp) is smaller and can drive stronger transgene expression than LAP1 (498 bp) after intravenous AAV-PHP.eB delivery,^13^ we assessed LAP2 in peripheral tissues and organs such as the liver, kidney, lung, and skeletal muscle. AAV provides the ability to drive tropism to specific tissues by using appropriate serotype and promoter combinations.^15^ AAV9 is a widely-used serotype with body-wide tropism,^16^ whereas AAV8 efficiently transduces liver and skeletal muscle.^17,18^ Recombinant AAV8 and AAV9 vectors share the ability to transduce peripheral tissues after systemic administration in mice. To date, both serotypes have routinely been used in clinical trials.^2^ To determine whether LAP2 drives efficient transgene expression in peripheral tissue and organ, AAV-LAP2 recombinants were packaged into both AAV8 and AAV9 capsids. The AAV constructs were delivered by systemic intravenous (retro-orbital; RO) and local (intramuscular; IM) routes of administration (ROA). Intravenous delivery is an attractive approach that is minimally invasive while providing widespread gene transfer and is a therapeutic option for clinical applications.^19,20^ Direct intramuscular administration can not only target the muscle, but also distant organs through the bloodstream.^21^ We performed comprehensive side-by-side studies using LAP2 and the strong and ubiquitous promoter human elongation factor-1 alpha (EF1α, 1264 bp). The tissue was examined 30 days post RO and IM administration in adult mice. We observed that AAV8- and AAV9-LAP2 drive potent transgene expression in liver, lung and kidney after RO injection, and in quadriceps (QUAD), tibialis anterior (TA), extensor digitorum longus (EDL), soleus (SOL) and gastrocnemius (GA) skeletal muscle after IM injection. In liver and skeletal muscle over 75% of cells showed LAP2-mCherry expression, while in kidney and lung over 40% of cells were mCherry positive. Next we compared the vector DNA biodistribution in liver, kidney, lung, and skeletal muscle. AAV8-LAP2 showed predominant tropism for liver after RO injection, with fewer copies found in kidney, lung, and skeletal muscle after RO and IM administration. In kidney, AAV9-LAP2 showed elevated level of biodistribution after RO delivery, whereas in lung and skeletal muscle, AAV8-LAP2 and AAV9-LAP2 showed levels of biodistribution that were significantly higher after IM injection. Abundant mCherry transcripts were detected in all tissue and organs confirming that LAP2 drives efficient transcription from the AAV DNA. Together these results demonstrate that after a single dose, AAV-LAP2 can drive widespread and potent transgene expression in peripheral tissue and organs comparable to that of the much larger EF1α promoter. These findings support the development of new therapies using LAP2 in cases where potent, broad, ubiquitous transgene expression is required.

## Results

### AAV8- and AAV9-LAP2 exhibit strong transcription and transgene expression in liver, kidney, and skeletal muscle after RO and IM routes of administration

AAV serotype and route of delivery may influence the transduction efficiency *in vivo*. In terms of specific transduction and tropism profile, it has been shown to be dependent on AAV capsid properties^22^. To characterize the performance of the LAP2 promoter in peripheral tissue and organs, we packaged the fluorescent reporter gene mCherry downstream of LAP2 into AAV8 and AAV9 capsids. We compared the efficiency of LAP2 with the ubiquitous EF1α promoter (Figure 1A). The AAV vectors were injected into mice at a dose of 5 × 10^11^ vector genomes per animal, either intravenously into the retro-orbital sinus or by local injection into the tibialis anterior (Figure 1B). Thirty days after injection, tissue was collected for histological and biochemical analysis (Figure 1C–1E).

**Figure 1.**
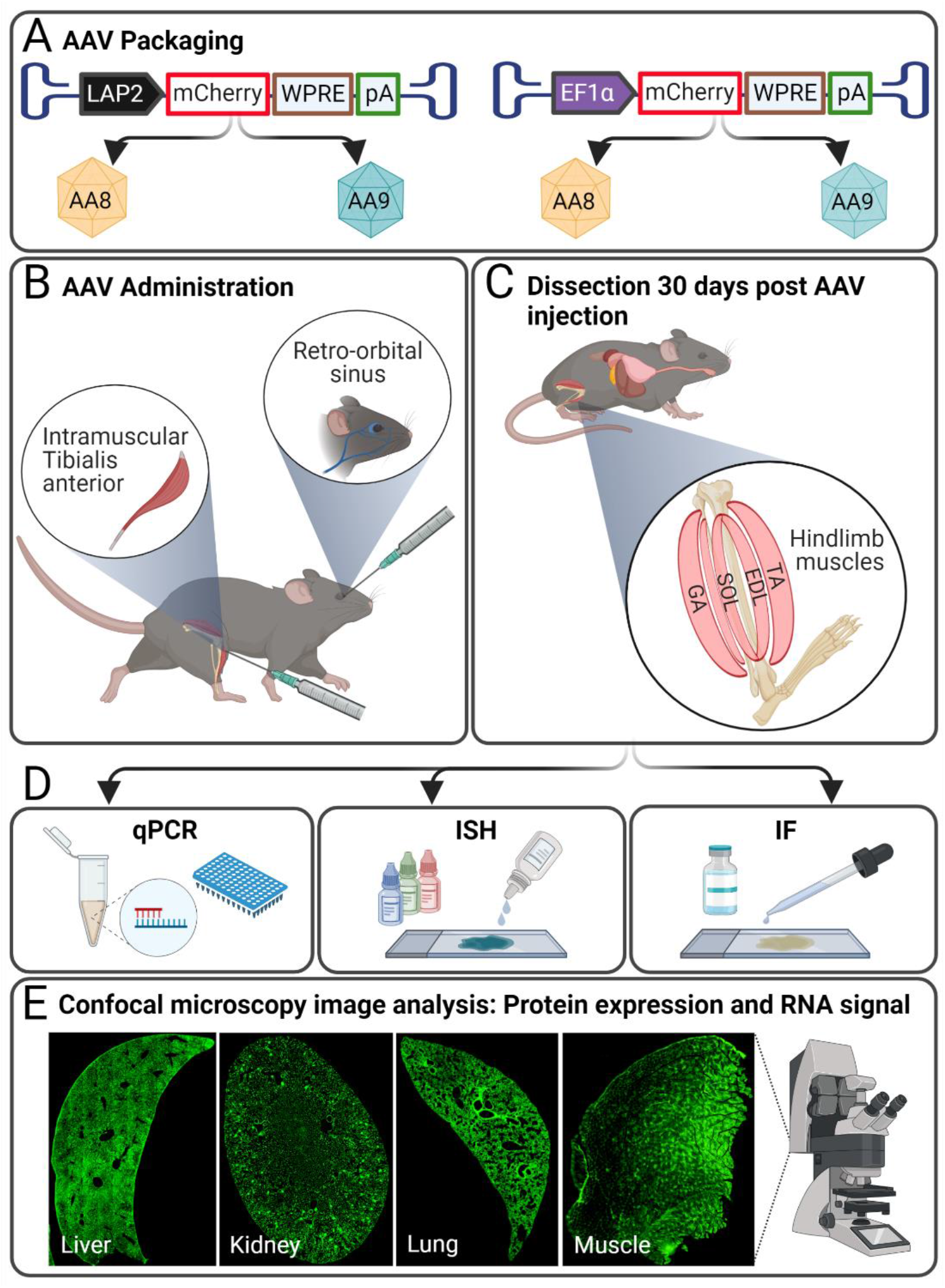
Constructs and workflow for analysis of the in vivo expression of AAV8- and AAV9-LAP2 vectors. (A) AAV packaging. The LAP2 and EF1α promoters were packaged into AAV8 and AAV9 vectors driving transcription of the mCherry fluorescent reporter. Expression cassette includes WPRE (woodchuck hepatitis virus post-transcriptional regulatory element) and SV40 polyA (simian virus 40 polyadenylation) polyadenylation site. B) AAV administration. Each AAV vector was administered either by unilateral intravenous injection into retro-orbital sinus or intramuscular injection at the tibialis anterior C57BL/6 mice at a dose 5 × 10^11^ vg per animal. C) Dissection and perfusion. Liver, kidney, lung, and skeletal muscles: quadriceps, tibialis anterior (TA), extensor digitorium longus (EDL), soleus (SOL), gastrocnemius (GA) was collected at 30 days post AAV injection. D) Experimental workflow. DNA from tissues was extracted after whole tissue homogenization for qPCR. Serial cross sections of the liver, kidney, and lung (20 µm thickness), and skeletal muscle (10 µm thickness) were cut using a cryostat for subsequent immunofluorescence (IF) and *in situ* hybridization (ISH). E) Confocal microscopy image analysis. Fluorescence imaging was conducted with a confocal microscope and analyzed by Image J and QuPath software. Representative images show phalloidin staining (green) for liver and kidney slices, and laminin staining (green) of lung and skeletal muscle. The image was created with biorender.com.

Overlay images of native mCherry expression (red), phalloidin fluorescent labelling of actin filaments to outline liver cells (green), and nuclei (blue) show that AAV8- and AAV9-LAP2 as well as AAV8- and AAV9-EF1α transduces hepatocytes near central veins (cv) (Figure 2A) and periportal regions (not shown) after intravenous and local administration. Similarly in kidney sections, phalloidin-stained cytoskeleton patterns (green) show that mCherry expression is restricted to the cortex and was enriched in the glomerulus (G), proximal tubule (PT), and distal convoluted tubule (DT) for both serotypes and promoters (Figure 2B). Both AAV8 and AAV9 vectors showed strong mCherry expression in liver and kidney after RO injection (Figure 2A-2B). In lung sections, laminin immunostaining (green) showed mCherry expression in alveolar cells (arrowhead) after delivery with AAV8- and AAV9-LAP2 by RO or IM injection (Figure 2C). There was no difference in the transgene expression between promoters. To analyze mCherry expression in skeletal muscle, we analyzed QUAD cross-sections stained with a laminin antibody (green) to define muscle fibers. In skeletal muscle mCherry had a mosaic pattern expression (asterisk) after IM administration with a comparable potency for both promoters (Figure 2D). We assessed the intensity of mCherry fluorescence driven by each test article in liver, kidney, lung, and skeletal muscle sections. As shown in Figure 2E1 and 2E2, fluorescence was significantly increased in liver and kidney after RO administration with AAV8- and AVV9-LAP2 compared to local IM injection (Table 1). However, there was no statistically significant difference between AAV capsid or promoters in liver (Figure 2E1; Table 1). Interestingly in kidney, AAV9-LAP2 and AAV9-EF1α showed significant differences compared to AAV8-LAP2 or AAV8-EF1α after IM injection (Figure 2E2; Table 1). Reporter fluorescence levels in lung were similar and not significantly different between AAV8- and AAV9-LAP2 after intravenous or local IM administration with both promoters (Figure 2E3; Table 1). In skeletal muscle, both AAV8- and AVV9-LAP2 showed significantly higher expression after local IM administration relative to intravenous administration but no significant differences between promoters or AAV serotype were determined (Figure 2E4; Table 1). Cell counts expressed as a percentage of mCherry-positive over total cells revealed significant differences in transduction efficiency between different routes of administration in liver (RO: ∼97% vs IM: ∼83%) for AAV8- and AAV9-LAP2 but showed no significant changes for AAV capsid or promoters (Figure 2F1; Table 1). In kidney, 55% of cells displayed mCherry expression after intravenous AAV9-LAP2 administration, and 40% of cells were mCherry positive for both AAV8-LAP2 by RO injection and AAV8- and AAV9-LAP2 by IM injection had fewer cells expressing mCherry (∼40%) (Figure 2F2; Table 1). This trend of ∼ 40% of total cells expressing mCherry was maintained in lung, after delivery with AAV8- and AAV9-LAP2 for both routes of delivery (Figure 2F3; Table 1). Conversely, the number cells expressing mCherry was significantly increased in the skeletal muscle (∼75%) after RO and IM administration of AAV8- and AAV9-LAP2 (Figure 2F4; Table 1). In terms of AAV serotype and promoters we did not observe significant differences in lung and skeletal muscle. Overall, these results demonstrate that the small LAP2 promoter drives strong and ubiquitous transgene expression, similar to the larger Ef1α promoter in peripheral tissue/organs with AAV8 or AAV9 capsids as determined by the route of administration.

**Table 1.** mCherry expression in mouse peripheral tissue.

| Tissue | Promoter | AAV serotype | Route of administration | Fluorescence Intensity (Mean ± SEM) | % mCherry Positive Cell (Mean ± SEM) |
| --- | --- | --- | --- | --- | --- |
| Liver | LAP2 | AAV8 | RO | 11645 ± 1056 | 97.5 ± 0.6 |
| | EF1α | AAV8 | RO | 11847 ± 1159 | 97.9 ± 0.7 |
| | LAP2 | AAV9 | RO | 11358 ± 765 | 94.2 ± 2.4 |
| | EF1α | AAV9 | RO | 9917 ± 614 | 98.6 ± 0.3 |
| | LAP2 | AAV8 | IM | 4854 ± 480 | 84.9 ± 2.4 |
| | EF1α | AAV8 | IM | 4144 ± 460 | 81.0 ± 2.2 |
| | LAP2 | AAV9 | IM | 5924 ± 692 | 83.7 ± 1.4 |
| | EF1α | AAV9 | IM | 5798 ± 985 | 82.8 ± 1.7 |
| Kidney | LAP2 | AAV8 | RO | 6026 ± 510 | 40.2 ± 1.2 |
| | EF1α | AAV8 | RO | 6010 ± 705 | 38.8 ± 1.8 |
| | LAP2 | AAV9 | RO | 6908 ± 209 | 54.7 ± 1.6 |
| | EF1α | AAV9 | RO | 6365 ± 361 | 40.6 ± 2.1 |
| | LAP2 | AAV8 | IM | 3288 ± 87 | 39.5 ± 2.3 |
| | EF1α | AAV8 | IM | 2569 ± 245 | 40.6 ± 2.3 |
| | LAP2 | AAV9 | IM | 4789 ± 211 | 42.1 ± 1.4 |
| | EF1α | AAV9 | IM | 4292 ± 175 | 41.4 ± 1.4 |
| Lung | LAP2 | AAV8 | RO | 906 ± 32 | 37.8 ± 1.7 |
| | EF1α | AAV8 | RO | 941 ± 53 | 40.9 ± 2.0 |
| | LAP2 | AAV9 | RO | 911 ± 32 | 40.3 ± 2.2 |
| | EF1α | AAV9 | RO | 796 ± 24 | 40.9 ± 2.1 |
| | LAP2 | AAV8 | IM | 908 ± 40 | 37.3 ± 1.7 |
| | EF1α | AAV8 | IM | 929 ± 56 | 42.3 ± 1.5 |
| | LAP2 | AAV9 | IM | 938 ± 33 | 39.2 ± 2.0 |
| | EF1α | AAV9 | IM | 827 ± 29 | 42.4 ± 2.5 |
| Skeletal muscle | LAP2 | AAV8 | RO | 3478 ± 198 | 75.4 ± 1.5 |
| | EF1α | AAV8 | RO | 2873 ± 462 | 78.1 ± 1.5 |
| | LAP2 | AAV9 | RO | 3803 ± 166 | 74.1 ± 1.3 |
| | EF1α | AAV9 | RO | 4008 ± 161 | 75.0 ± 1.3 |
| | LAP2 | AAV8 | IM | 4560 ± 167 | 76.3 ± 1.5 |
| | EF1α | AAV8 | IM | 4293 ± 221 | 76.7 ± 1.3 |
| | LAP2 | AAV9 | IM | 5113 ± 167 | 76.0 ± 1.7 |
| | EF1α | AAV9 | IM | 5208 ± 279 | 74.7 ± 1.5 |

RO, retro-orbital injection; IM, intramuscular injection.

**Figure 2.**
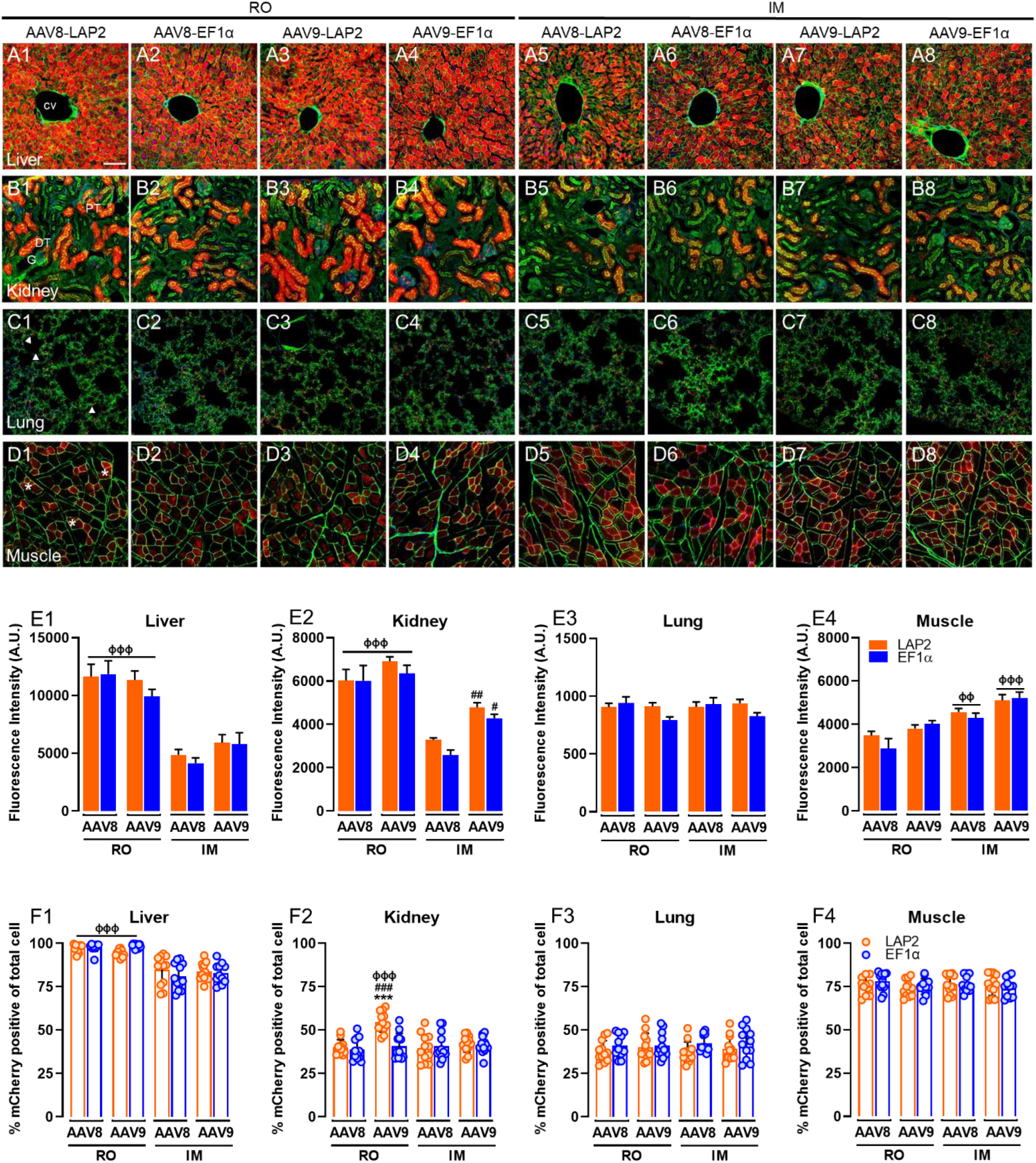
Characterization of LAP2 promoter after AAV8 and AAV9 intravenous and intramuscular administration. Representative confocal images show native mCherry fluorescence (red) A) around to central vein (cv) in liver, B) glomerulus (G), proximal tubule (PT), and distal convoluted tubules (DT) in kidney, C) alveolar cells (arrowhead) in lung, and D) myofiber (asterisks) in skeletal muscle after 30 days post administration. Liver and kidney sections were immunostained with phalloidin (green) and lung and skeletal muscle, with laminin (green) counterstained with DAPI (blue). mCherry expression after delivery with AAV8-LAP2 and AAV9-LAP2 by (A1-A4, B1-B4, C1-C4, D1-D4) RO injection and (A5-A8, B5-B8, C5-C8, D5-D8) IM injection is compared side-by-side with EF1α promoter in liver, kidney, lung, and skeletal muscle respectively. Scale bar, 100 µm. (E) Quantification of fluorescence intensity of native mCherry driven by AAV8-LAP2 and AAV9-LAP2 versus AAV8-EF1α and AAV9-EF1α at 30 days post administration by RO and IM injection is shown in (E1) liver, (E2) kidney, (E3) lung, and (E4) skeletal muscle. (F) Quantification of percentage of mCherry positive of total cells driven by AAV8-LAP2 and AAV9-LAP2 versus AAV8-EF1α and AAV9-EF1α at 30 days post administration by RO and IM injection is shown in (F1) liver, (F2) kidney, (F3) lung, and (F4) skeletal muscle. Data are reported as mean ± SEM; n = 3 animals per condition (four tissue section were analyzed for each animal). ^***^p < 0.001 compared to promoters. ^#^p < 0.033, ^##^p < 0.002, ^###^p<0.001 compared to serotype. ^ϕϕϕ^p < 0.001 compared to route of administration. Two-way ANOVA with Turkey’s post hoc test. (RO) retro-orbital injection; (IM) intramuscular injection.

### AAV8- and AAV9-LAP2 lead to widespread biodistribution and transgene expression that is influenced by the route of administration

To determine whether there are differences in the biodistribution of AAV8-LAP2 and AAV9-LAP2, we quantified the vector genome copy number per cell in peripheral tissue and organs. AAV8-LAP2 showed the highest copy number in liver (102 VGCN/cell) compared to AAV9-LAP2 (61 VGCN/cell), both after RO injection and AAV8- and AAV9-LAP2 after IM injection (average 24 VGCN/cell) (Figure 3A; Table 2). In the kidney, which was orders of magnitude lower than the liver, AAV9-LAP2 by RO injection showed elevated level of biodistribution (1.3 VGCN/cell) in contrast to AAV9-LAP2 (0.5 VGCN/cell) by IM injection (Figure 3B; Table 2). There was no significant VGCN/cell difference for AAV8 and AAV9 regardless of route. In the lung, AAV9-LAP2 delivery after IM injection resulted in significantly higher biodistribution (5.9 VGCN/cell) than AAV8-LAP2 by same ROA (3.6 VGCN/cell) and AAV8- and AAV9-LAP2 by RO injection (average 1.4 VGCN/cell) (Figure 3C; Table 2). In skeletal muscle, AAV8- and AAV9-LAP2 by IM administration (average 7.3 VGCN/cell) showed levels of biodistribution that were significantly higher than AAV8- and AAV9-LAP2 following RO injection (average 1.3 VGCN/cell) (Figure 3D; Table 2). In terms of VGCN between promoters no significant differences were observed in the different tissues.

**Table 2.** Vector genome copies number (VGCN) per cell in mouse peripheral tissue.

| Tissue | Promoter | AAV serotype | Route of administration | VGCN/cell<br>(Mean ± SEM) |
| --- | --- | --- | --- | --- |
| Liver | LAP2 | AAV8 | RO | 101.9 ± 20.9 |
| | EF1α | AAV8 | RO | 64.6 ± 14.0 |
| | LAP2 | AAV9 | RO | 61.4 ± 14.7 |
| | EF1α | AAV9 | RO | 56.7 ± 9.1 |
| | Buffer | | RO | 3.0 ± 0.8 |
| | LAP2 | AAV8 | IM | 29.1 ± 5.8 |
| | EF1α | AAV8 | IM | 18.0 ± 4.5 |
| | LAP2 | AAV9 | IM | 18.2 ± 4.9 |
| | EF1α | AAV9 | IM | 17.2 ± 1.2 |
| | Buffer | | IM | 0.7 ± 0.5 |
| Kidney | LAP2 | AAV8 | RO | 0.8 ± 0.2 |
| | EF1α | AAV8 | RO | 0.7 ± 0.3 |
| | LAP2 | AAV9 | RO | 1.3 ± 0.3 |
| | EF1α | AAV9 | RO | 1.2 ± 0.2 |
| | Buffer | | RO | 0.2 ± 0.0 |
| | LAP2 | AAV8 | IM | 0.5 ± 0.1 |
| | EF1α | AAV8 | IM | 0.4 ± 0.1 |
| | LAP2 | AAV9 | IM | 0.5 ± 0.1 |
| | EF1α | AAV9 | IM | 0.7 ± 0.2 |
| | Buffer | | IM | 0.0 ± 0.0 |
| Lung | LAP2 | AAV8 | RO | 1.4 ± 0.3 |
| | EF1α | AAV8 | RO | 1.5 ± 0.7 |
| | LAP2 | AAV9 | RO | 1.4 ± 0.1 |
| | EF1α | AAV9 | RO | 1.2 ± 0.4 |
| | Buffer | | RO | 0.1 ± 0.0 |
| | LAP2 | AAV8 | IM | 3.6 ± 1.2 |
| | EF1α | AAV8 | IM | 1.8 ± 0.6 |
| | LAP2 | AAV9 | IM | 5.9 ± 1.3 |
| | EF1α | AAV9 | IM | 4.7 ± 1.0 |
| | Buffer | | IM | 0.1 ± 0.0 |
| Skeletal muscle | LAP2 | AAV8 | RO | 1.4 ± 0.4 |
| | EF1α | AAV8 | RO | 1.5 ± 0.8 |
| | LAP2 | AAV9 | RO | 1.1 ± 0.2 |
| | EF1α | AAV9 | RO | 1.4 ± 0.4 |
| | Buffer | | RO | 0.3 ± 0.1 |
| | LAP2 | AAV8 | IM | 7.3 ± 1.2 |
| | EF1α | AAV8 | IM | 4.7 ± 0.8 |
| | LAP2 | AAV9 | IM | 7.2 ± 1.5 |
| | EF1α | AAV9 | IM | 5.8 ± 1.0 |
| | Buffer | | IM | 0.1 ± 0.0 |

RO, retro-orbital injection; IM, intramuscular injection.

**Figure 3.**
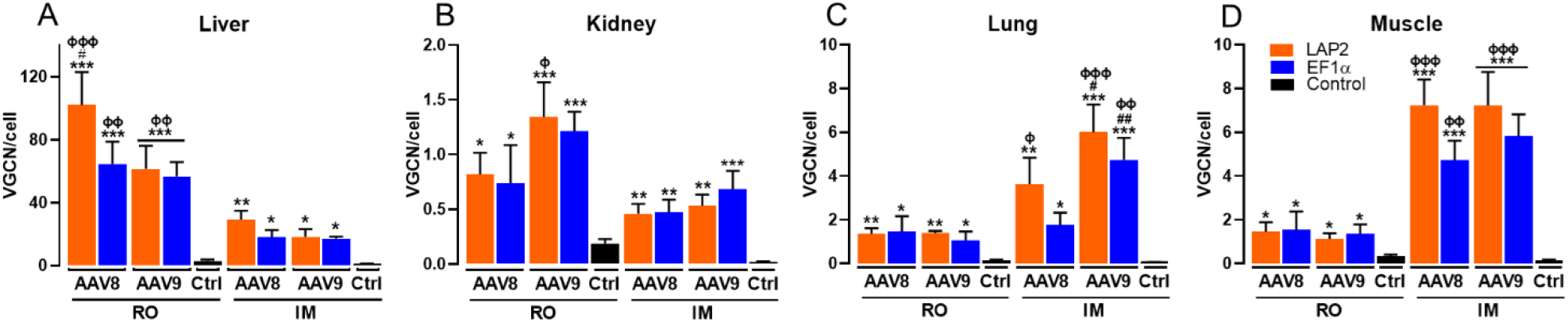
AAV8- and AAV9-LAP2 vectors DNA biodistribution in transduced peripheral tissue. Vector genome copies number (VGCN) per cell in A) liver, B) kidney, C) lung, and D) skeletal muscle was evaluated using qPCR at 30 days post administration. Analysis of vector biodistribution after RO or IM injection of AAV8-LAP2 and AAV9-LAP2 versus AAV8-EF1α and AAV9-EF1α. Data are reported as mean ± SEM; n = 3 animals per condition. *p < 0.033, ^**^p < 0.002, ^***^p < 0.001 compared to control (Ctrl). ^#^p < 0.033, ^##^p < 0.002 compared to serotype. ^ϕ^p < 0.033, ^ϕϕ^p < 0.002, ^ϕϕϕ^p < 0.001 compared to route of administration. Two-way ANOVA with Turkey’s post hoc test. (RO) retro-orbital injection; (IM) intramuscular injection.

Additionally, we measured mCherry transcription by fluorescent RNA *in situ* hybridization. The amount of mCherry mRNA was measured in liver, kidney, lung, and skeletal muscle after AAV8- and AAV9-LAP2 RO and IM administration (Figure 4A-4D). The presence of mCherry messenger RNA demonstrates persistent LAP2-driven transcription. Altogether the findings demonstrate that AAV-LAP2 has a tropism and expression profile equivalent to that of the larger EF1α promoter.

**Figure 4.**
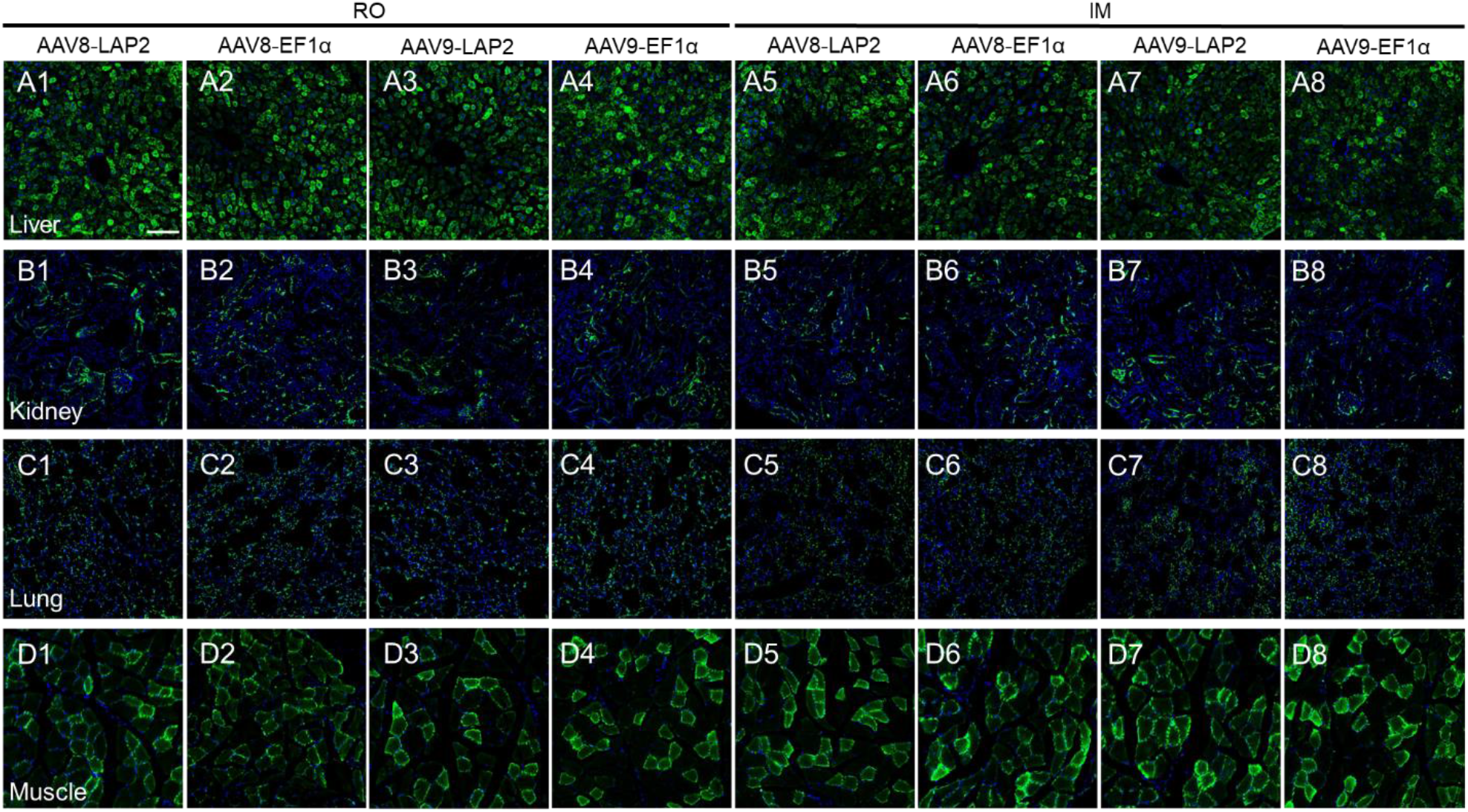
AAV8- and AAV9-LAP2-mCherry transcription in liver, kidney, skeletal muscle, and lung tissue after RO and IM administration measured by ISH. *In situ* hybridization was performed on A) liver, B) kidney, C) lung, and D) skeletal muscle samples using the RNAscope Multiplex Assay. Probe hybridized to RNA were imaged with a confocal microscope. Transgene RNA (green) and nuclear counterstain, DAPI (blue). Transgene transcription after delivery with AAV8-LAP2 and AAV9-LAP2 by (A1-A4, B1-B4, C1-C4, D1-D4) RO injection and (A5-A8, B5-B8, C5-C8, D5-D8) IM injection in side-by-side comparison with EF1α promoter in liver, kidney, lung, and skeletal muscle, respectively. Scale bar, 100 µm. (RO) retro-orbital injection; (IM) intramuscular injection.

### Intramuscular AAV8- and AAV9-LAP2 administration leads to potent transgene expression in type 2 (fast-twitch muscle) fibers

To compare transduction efficacy observed in QUAD muscle (Figure 2D, 2E4, 2F4; Table 1) with others skeletal muscles after AAV8- and AVV9-LAP2 by IM injection, we quantified mCherry expression in hindlimb skeletal muscles: TA, EDL, SOL, and GA. The mCherry fluorescent intensity measured from the AAV8- and AAV9-LAP2 after IM administration was significantly higher than that of RO administration in TA, EDL, and SOL (Figure 5A1-5A3; Table 3). In GA, AAV8- and AVV9-LAP2 do not show significant differences between and AAV serotype and ROA, except for AAV9-EF1α after RO and IM delivery (Figure 5A4; Table 3). Conversely, the number of cells expressing mCherry after with AAV8- and AAV9-LAP2 delivery had higher efficacy (∼75%) without differences between ROA in TA, EDL, and GA (Figure 5B1, 5B2, 5B4; Table 3). Despite the lower mCherry expression per cell in the SOL there was no significative differences between RO and IM injection (Figure 5B3; Table 3). In terms of AAV serotype and promoters we did not observe significant differences between transduced skeletal muscles.

**Table 3.** mCherry expression in mouse hindlimb skeletal muscle.

| Hindlimb muscle | Promoter | AAV serotype | Route of administration | Fluorescence Intensity (Mean ± SEM) | % mCherry Positive Cell (Mean ± SEM) |
| --- | --- | --- | --- | --- | --- |
| TA | LAP2 | AAV8 | RO | 2681 ± 157 | 72.4 ± 1.5 |
| | EF1α | AAV8 | RO | 2429 ± 198 | 71.4 ± 2.1 |
| | LAP2 | AAV9 | RO | 3001 ± 203 | 78.8 ± 1.6 |
| | EF1α | AAV9 | RO | 2883 ± 309 | 76.5 ± 2.4 |
| | LAP2 | AAV8 | IM | 3576 ± 221 | 81.9 ± 1.4 |
| | EF1α | AAV8 | IM | 3249 ± 240 | 75.2 ± 2.7 |
| | LAP2 | AAV9 | IM | 3710 ± 231 | 76.6 ± 1.9 |
| | EF1α | AAV9 | IM | 3329 ± 215 | 79.3 ± 1.6 |
| EDL | LAP2 | AAV8 | RO | 2206 ± 261 | 78.1 ± 1.5 |
| | EF1α | AAV8 | RO | 2389 ± 370 | 77.1 ± 1.9 |
| | LAP2 | AAV9 | RO | 2493 ± 253 | 79.3 ± 1.7 |
| | EF1α | AAV9 | RO | 2595 ± 368 | 72.5 ± 1.9 |
| | LAP2 | AAV8 | IM | 3295 ± 142 | 78.9 ± 1.6 |
| | EF1α | AAV8 | IM | 2879 ± 166 | 77.3 ± 1.9 |
| | LAP2 | AAV9 | IM | 3489 ± 159 | 76.6 ± 2.0 |
| | EF1α | AAV9 | IM | 3571 ± 172 | 79.3 ± 1.7 |
| SOL | LAP2 | AAV8 | RO | 2751 ± 185 | 60.2 ± 1.3 |
| | EF1α | AAV8 | RO | 2346 ± 217 | 59.9 ± 0.9 |
| | LAP2 | AAV9 | RO | 3157 ± 264 | 59.1 ± 1.3 |
| | EF1α | AAV9 | RO | 2961 ± 327 | 61.1 ± 1.1 |
| | LAP2 | AAV8 | IM | 3687 ± 88 | 61.2 ± 1.8 |
| | EF1α | AAV8 | IM | 3422 ± 173 | 62.2 ± 1.8 |
| | LAP2 | AAV9 | IM | 4539 ± 110 | 62.6 ± 1.7 |
| | EF1α | AAV9 | IM | 3976 ± 135 | 63.5 ± 2.1 |
| GA | LAP2 | AAV8 | RO | 2867 ± 138 | 71.8 ± 1.6 |
| | EF1α | AAV8 | RO | 2960 ± 115 | 71.7 ± 1.5 |
| | LAP2 | AAV9 | RO | 3113 ± 243 | 78.5 ± 1.2 |
| | EF1α | AAV9 | RO | 3069 ± 148 | 77.4 ± 1.5 |
| | LAP2 | AAV8 | IM | 3357 ± 306 | 78.1 ± 1.4 |
| | EF1α | AAV8 | IM | 3220 ± 102 | 78.9 ± 1.7 |
| | LAP2 | AAV9 | IM | 3569 ± 193 | 79.9 ± 1.4 |
| | EF1α | AAV9 | IM | 3797 ± 158 | 80.1 ± 1.1 |

TA, tibialis anterior; EDL, extensor digitorum longus; SOL, soleus; GA, gastrocnemius; RO, retro-orbital injection; IM, intramuscular injection.

**Figure 5.**
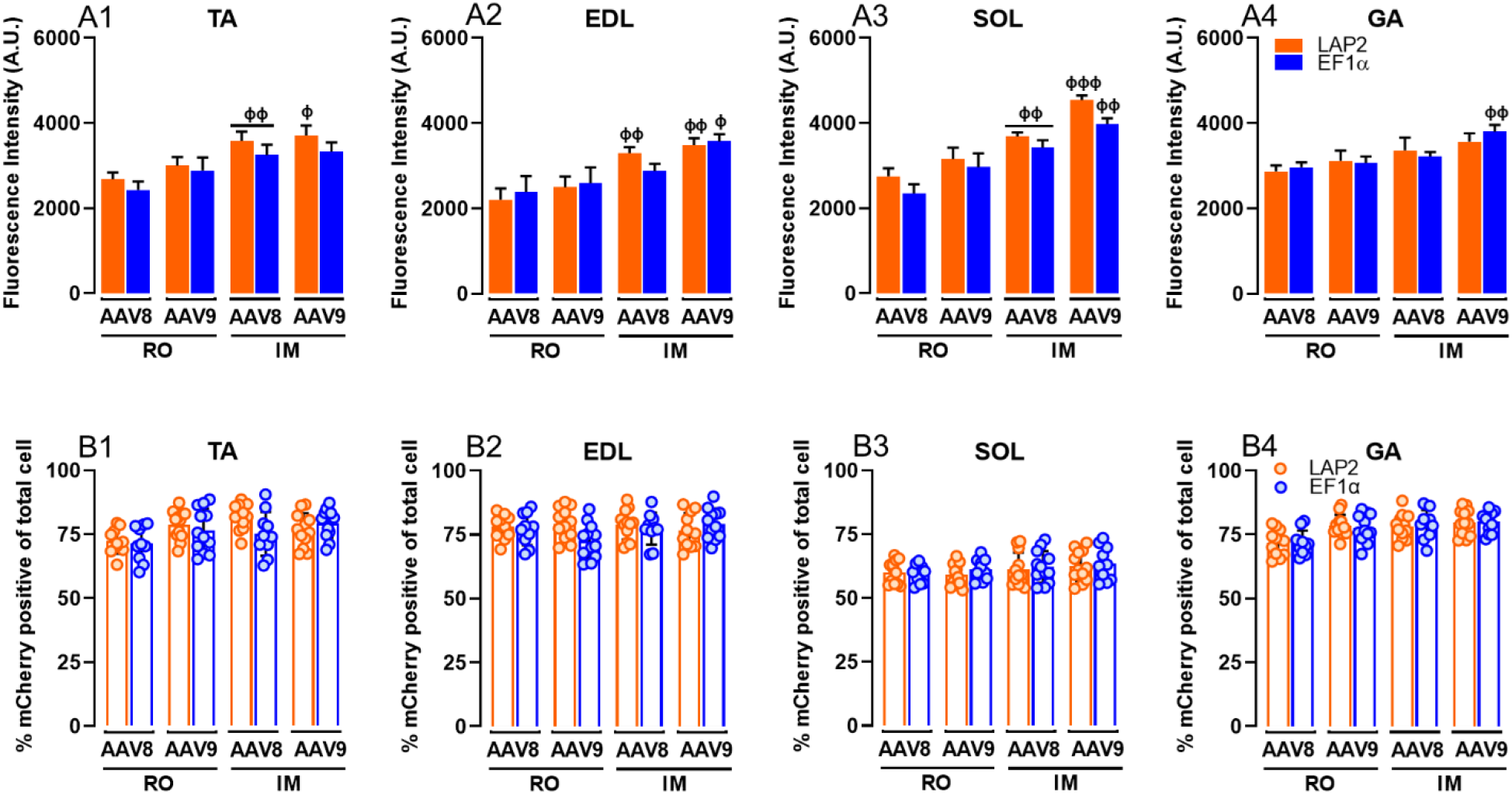
Comparison of mCherry expression of AAV8- and AAV9-LAP2 transduction profiles following RO and IM administration in hindlimb skeletal muscle. (A) Quantification of fluorescence intensity of native mCherry driven by AAV8-LAP2 and AAV9-LAP2 versus AAV8-EF1α and AAV9-EF1α at 30 days post administration by RO and IM injection is shown in (A1) TA, (E2) EDL, (A3) SOL, and (A4) GA. (B) Quantification of percentage of mCherry positive of total cells driven by AAV8-LAP2 and AAV9-LAP2 versus AAV8-EF1α and AAV9-EF1α at 30 days post administration by RO and IM injection is shown in (B1) TA, (B2) EDL, (B3) SOL, and (B4) GA. Data are reported as mean ± SEM; n = 3 animals per condition (four tissue section were analyzed for each animal). ^ϕ^p < 0.033, ^ϕϕ^p < 0.002, ^ϕϕϕ^p < 0.001 compared to route of administration. Two-way ANOVA with Turkey’s post hoc test. (TA) tibialis anterior; (EDL) extensor digitorum longus; (SOL) soleus; (GA) gastrocnemius; (RO) retro-orbital injection; (IM) intramuscular injection.

Since we detected a mosaic mCherry expression pattern for both AAV8- and AVV9-LAP2 (Figure 6), we analyzed whether the LAP2 promoter was driving a fiber type preference after IM administration. We performed immunostaining with anti-MyHC-2 (myosin heavy chain type 2) in the QUAD and hindlimb skeletal muscles. Overlay images for native mCherry expression (red), laminin/MyHC-2 (green), and nuclei (blue) show that the majority of the mCherry-positive correspond to myofibers type 2 (fast-twitch muscle) in TA, EDL, GA, and QUAD (Figure 6A, 6B, 6D, 6E; arrows) in contrast to SOL, which is rich in slow-twitch muscle (Figure 6C). The cells expressing mCherry that were not co-labelled with the MyHC-2 signal might correspond to other unidentified MyHC types (type 1, 4, 7)^23^ (Figure 6A, 6B, 6C, 6D, 6E; arrowheads). These results demonstrate that both AAV8- and AVV9-LAP2 drive efficient and potent transgene expression preferentially in fast-twitch muscle.

**Figure 6.**
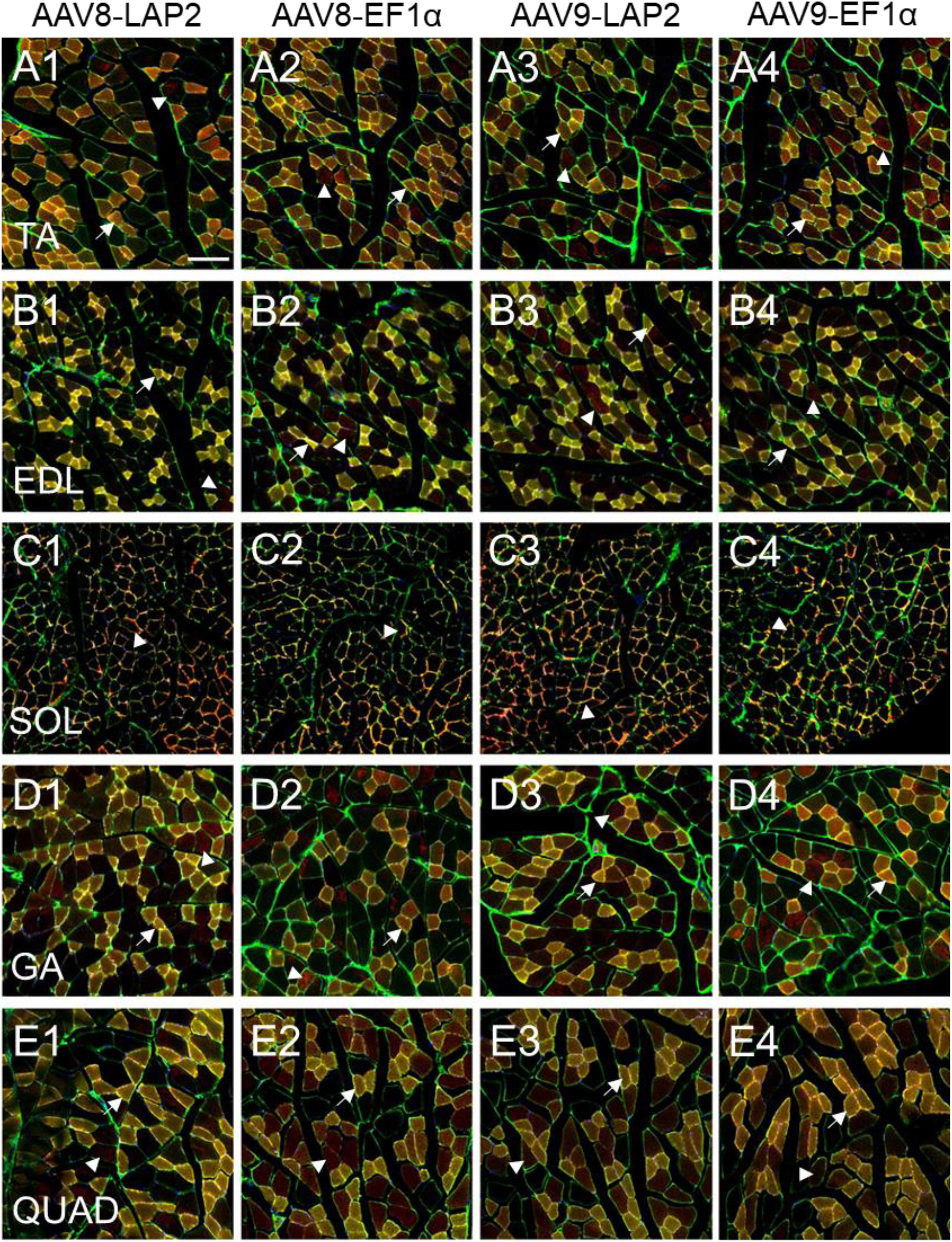
Intramuscular AAV8-LAP2 and AAV9-LAP2 administration leads to potent transgene expression in type 2 (fast-twitch muscle) fibers. Representative confocal images show immunofluorescence of skeletal muscle sections after 30 days post administration. Native mCherry (red) and muscle fiber outline (laminin)/type 2 fiber (MyHC-2) (green) was counterstained with DAPI (blue). Transgene expression after delivery with AAV8-LAP2 and AAV9-LAP2 by IM injection in side-by-side comparison with EF1α promoter in (A) TA, (B) EDL, (C) SOL, (D) GA and (E) QUAD, respectively. Example of matching of type 2 fiber and mCherry expression are marked with arrow, and not merge, with arrowheads. Scale bar, 100 µm. (TA) tibialis anterior; (EDL) extensor digitorum longus; (SOL) soleus; (GA) gastrocnemius; (QUAD) quadricep.

## Discussion

Gene therapy using AAV for gene transfer is a relatively novel and promising arm of translational medicine.^24,25^ The application of AAV-based gene therapy shows potential for a wide spectrum of diseases.^26^ The efficacy of such therapies depend on the potent and persistent expression of the transgene into the target cell and affected tissue and organs. Gene transfer and expression can be optimized according to the specific disease paradigm and it is determined by the AAV serotype, dose, route of administration and regulatory elements such as promoters.^27^ Although different AAV capsids can transduce a plethora of organs, it has been demonstrated that the route of administration can significantly influence the transduction efficiency and biodistribution profile of AAV vectors.^21,28,29^ Further, effective gene transfer can be achieved by using strong promoters. Ubiquitous promoters provide the trade-off of being typically more potent than tissue-specific promoters.^7^ Some clinical trials use ubiquitous and strong promoters such as CBh or miniCBA. However, these promoters are susceptible to transcriptional silencing that would render them incompatible with single-dose, “one and done” therapies.^30^ Constitutive promoters like cytomegalovirus (CMV, 800 bp), CMV enhancer fused to the chicken beta-actin promoter (CAG, 1.7 kb), or EF1α provide efficient and long-lasting transgene expression, but the large size limit the genetic payload available for the therapeutic transgene.^8,13^ Thus, the availability of small, strong and durable gene promoters is paramount for successful gene therapies. Here, we report a short, potent, constitutive, and persistent promoter, called LAP2. This promoter has been described to be resistant to epigenetic silencing during latency, chronically driving the transcription of LATs in neurons.^12^ We previously demonstrated that AAV-LAP2 acts as a panneuronal promoter, with a long-lasting and potent transgene expression profile in the brain and spinal cord.^13^ However, earlier studies suggest that LAP would also have a determining role in the infectivity of PRV in peripheral organs,^14^ and especially be responsible for the production of LATs in both neuronal and non-neuronal cells.^10,11,31^ Thus, we determined whether LAP2 drives transgene expression in peripheral tissue/organs and did a side-by-side comparison with the ubiquitous promoter EF1α.

To date, there is a significant number of clinical trials using AAV8 and AAV9 since they have a broader tropism than other natural serotypes.^32^ Excluding CNS and ophthalmological diseases, these AAV vectors can be directly administered to patients by intravenous, IM or site-specific increasing the spectrum of treatments with gene therapy.^2^ Thus, we generated AAV8- and AAV9-LAP2 constructs, and performed comprehensive side-by-side experiments after 30 days post RO and IM administration in adult mice. We observed efficient by AAV8- and AAV9-LAP2 transduction in liver, kidney, and skeletal muscle, and weak transgene expression in lung after IM and RO injection. The levels of mCherry expression were significantly higher in hepatocytes with predominant pericentral and periportal transduction after intravenous AAV8- and AAV9-LAP2 administration. Previous studies have shown that the patterns of AAV transgene expression along a portocentral axis, called liver zonation, vary among different species.^33^ When liver-specific thyroid hormone binding globulin (TBG; 800 bp), CB and CMV promoters are administered intravenously, transgene expression is mainly pericentral in dogs, while in non-human primates is periportal^34^. These findings suggest that the promoters do not discriminate between liver zonation.^34,35^. Notably, livers from newborn or infant mice and rhesus monkeys show a more uniform transgene expression distribution, with no predilection for central or periportal venous areas after intravenous AAV8 delivery.^33,34^ Our results are particularly relevant for liver diseases, where early metabolic dysfunction occurs along the axis of the hepatic lobule. Moreover, AAV8-LAP2 vector genome copies after RO administration revealed an order of magnitude higher than that of other tissue and organs. As expected, abundant mCherry transcript was detected after RO administration for both AAV8- and AAV9-LAP2. Gene therapy has been developed for liver monogenic disorders with secreted protein such as hemophilia A (deficiency in factor VIII, FVIII), hemophilia B (deficiency in factor IX, FIX), and α1-antitripsyn deficiency (AAT deficiency, an inherited condition that raises the risk for lung and liver disease).^36^ Although, the coding sequences of these enzymes have been shortened,^37^ the newer versions remain at the payload limit of AAV vectors, further demonstrating the unmet and immediate need of small, potent and persistent promoters. In this scenario, our findings show that LAP2, with only 404 bp, drive widespread, potent, transgene expression in liver, influenced by AAV serotype and route of administration.

In renal histochemical analyses, the luminal border of the PT cells shows intense phalloidin fluorescence merging with mCherry expression. While the DT cells with only a thin bundle of actin filaments lining the luminal side display lower levels of mCherry-like glomerular regions. In contrast, the collecting duct in the cortex and transition zone between the outer and inner medullary regions did not show mCherry expression. Side-by-side comparative results show that AAV9-LAP2 provided strong transgene expression in renal cortex after RO delivery compared to IM injection. We also demonstrated that biodistribution as vector genomes per cell after intravenous delivery was significantly higher for AAV9-LAP2 than AAV9-EF1α. Moreover, the levels of mCherry transcript after RO delivery with AAV8- and AAV9-LAP2 are like what we observed for IM administration. These results are consistent with the findings of previous studies showing that retrograde infusion through the ureter of AAV9-CMV, display efficient transgene expression to PT, DT and collecting duct cells^38,39^. The kidney is composed of several segments, each with different functions attributed to the expression of single genes that regulate fluid and electrolyte homeostasis^40^. It has been demonstrated that naturally occurring AAV serotypes can transduce specific kidney segments with low level efficiency following intravenous injection^41,42^. In this scenario, the use of a strong promoter is essential to achieve sufficient transgene expression. However, kidney-specific promoters such as kidney-specific cadherin (KSPC; 1.3 kb), Na^+^/glucose co-transporter (SGLT2; 1.1 kb), and E-cadherin (ECAD; 1.25 kb), are typically large in size^43^. Our results demonstrate that transgenic expression driven by the small and potent LAP2 promoter in specific segments of the renal nephron, is useful both for basic research as well as for clinical trials.

In this study, both AAV serotypes and routes of administration evaluated, yielded similar transduction efficiency and transgene expression in lungs. Low mCherry levels and similar mCherry RNA levels were observed for AAV8- and AAV9-LAP2 after RO and IM routes of administration. Surprisingly, AAV-LAP2 following IM administration, showed genome copies per cell that were significantly higher than those measured for AAV-LAP2 after intravenous administration. In the context of RO and IM routes, it has been demonstrated that natural AAV as well as new engineered capsids, failed to generate efficient lung transduction^42^. Preclinical alpha-1-antitrypsin (AAT) deficiency studies using AAV1-CMV vector following IM administration with doses up 10^13^ vg, demonstrated dose-dependent AAT serum levels that peaked 30 days after vector administration and declined after 90 days^44^. Recent studies have been focused on improvements to AAV capsid to induce rapid AAV trafficking, entry to the nucleus and intracellular processing, avoiding early ubiquitination and degradation^45^. There are challenges in developing effective therapies in the lung due to anatomical differences between mouse and humans, and differences in the types of receptors expressed in the AAV capsid^28^. The LAP2 promoter could improve AAV vectors targeting the lung by lowering the dose while increasing transgene expression levels. Furthermore, LAP2 could be optimal for lung studies that require moderate to low transgene expression.

Consistent with previous studies, we show that high-efficiency transduction of LAP2 in skeletal muscle results in a mosaic pattern,^46^ that might be explained by the preferential mCherry expression in myofibers regardless of AAV-LAP2 serotype and route of administration. Studies with AAV9-CMV have shown higher transduction of TA muscle compared to the SOL muscle, suggesting that not all myofiber types are equally permissive to AAV vectors.^47^ Furthermore, previous studies suggest that CMV promoters do not discriminate between myofiber types,^48,49^ given that SOL muscle consists of mainly type 1 (slow-twitch) fibers, whereas TA, EDL, GA and QUAD muscle have more type 2 (fast-twitch) fibers.^50^ Consistent with our results, we observed that after IM administration of AAV8- and AAV9-LAP2, the fluorescence mCherry intensity and DNA biodistribution increased compared to that of intravenous administration. However, the levels of mCherry transcript and protein were the same for both ROA and AAV8- and AAV9-LAP2 showed efficient tissue transduction. These findings hold great promise for the treatment of several skeletal muscle diseases, such as Duchenne muscular dystrophy and Pompe disease requiring muscle gene transfer with AAV8 and AAV9 vectors.^21^ These results show that AAV-LAP2 drive efficient transduction following RO and IM administration of AAV8 and AAV9 with preferential mCherry expression in fast-twitch myofiber expressing MyHC-2 isotype.

In summary, we provide compelling evidence to support that the LAP2 promoter can drive strong, persistent, and widespread transgene expression in peripheral tissue and organs including liver, kidney, and skeletal muscle, and mild transgene expression in lung after IM and RO routes of administration. A direct side-by-side comparison between LAP2 and EF1α, demonstrates that regardless of the AAV serotype and route of administration, LAP2 is as powerful as EF1α promoter despite being 66% smaller in size, overcoming the limitations of the AAV vector.

## Materials and Methods

### Construction and Production of AAVs

AAV plasmids were produced as previously described.^13^ Each AAV plasmids were packaged into AAV8 and AAV9 serotype. The AAV titer was measured by qPCR using TaqMan (Thermo Fisher Scientific, Rockford, IL, USA) and reported as genome copies (GC)/ml. All AAV vectors were produced by the PNI Viral Core Facility (Princeton Neuroscience Institute, Princeton University). AAV plasmids contain mCherry driven by the LAP2 or EF1α promoter and include WPRE (woodchuck hepatitis virus post-transcriptional regulatory element) and SV40 polyA signal (simian virus 40 polyadenylation).

### Animals

Five-week-old male C57BL/6J mice were obtained from The Jackson Laboratory (Bar Harbor, ME, USA) and kept on a 12h light/dark cycle (lights on at 7:00 am) with ad libitum food and water. Special care was taken to minimize suffering and to reduce the number of animals used to the minimum required for statistical inference. The Princeton University Institutional Animal Care and Use Committee (protocols 1947-19) approved protocols.

### Vector administration

Unilateral retro-orbital sinus administration in adult mice was performed as previously described.^13^ Animals previously anesthetized received a total of 5 × 10^11^ vg of AAV in a volume of 100 µl. For intramuscular injection, unilateral TA muscle was injected with AAV according to established method.^51^ Right TA muscle received a total of 5 × 10^11^ vg of AAV in a volume of 50 µl.

### Tissue Sample Processing

Thirty days post-injection, mice were deeply anesthetized with ketamine (400 mg/kg)/xylazine (50 mg/kg) by intraperitoneal injection and then perfused with 4% paraformaldehyde in 0.1 M phosphate buffer saline (PBS) (Fisher Scientific, Waltham, MA, USA). Liver, kidney, lung, and skeletal muscle were dissected and then fixed in 4% paraformaldehyde in PBS for 1 day. Different skeletal muscle: TA, EDL, SOL, GA, and QUAD, were dissected to established method.^23^ Organs/tissue were then serially incubated in 15% and 30% sucrose in PBS overnight at 4°C. Tissue was placed in an embedding mold (Sigma-Aldrich, The Woodlands, TX, USA) with OCT (Tissue-Tek, Torrance, CA, USA), frozen in dry ice, and stored at -80°C until cryosection. Serial cross sections of the liver, kidney, and lung (20 µm thickness), and skeletal muscle (10 µm thickness) were cut using a Leica CM3050 S cryostat (Leica Biosystems, Buffalo Grove, IL, USA).

### Immunostaining

Cryosections mounted on a glass slide (Fisher Scientific, Waltham, MA, USA) were blocked with 3% bovine serum albumin (BSA), 2% donkey serum, and 0.5% Triton X-100 (Sigma-Aldrich, St. Louis, MO, USA) for 1 h. The sections were incubated with primary antibodies overnight at 4°C and further incubated with appropriate secondary antibodies for 1 h at RT. Skeletal muscle were stained with Myosin 4 monoclonal antibody (MF20, MyHC-2) (1:200; Thermo Fisher Scientific, Rockford, IL, USA) and Laminin gamma 1 Antibody (A5) (1:400; Novus Biologicals, Centennial, CO, USA) with secondary antibody, Alexa Fluor 488 donkey anti-mouse IgG and Alexa Fluor 647 donkey-rat IgG (1:1,000; Thermo Fisher Scientific, Rockford, IL, USA). Liver, kidney, and lung cryosections after blocking were stained with a 1:200 dilution of Alexa Fluor 488 Phalloidin (Thermo Fisher Scientific, Rockford, IL, USA) for 1 h. Cell nuclei were counterstained with 0.5 mg/mL DAPI for 5 min (Thermo Fisher Scientific, Rockford, IL, USA) and mounted using Vectashield Vibrance antifade mounting media (Molecular Probes, Eugene, OR, USA).

### Imaging and Analysis

Imaging was conducted with a Leica SP8-LSCM confocal microscope (Leica Microsystems, Wetzlar, Germany) using a 20X objective and 0.5 μm z steps. Z-stacks were generated with the ImageJ software. Four random image areas were selected for each tissue sections collected per animal (n=3). Image quantifications were analyzed using QuPath 0.3.0 software.^52^ The algorithm for DAPI detection was based on a pixel classifier and was directed to automatically identify the islet areas (region of interest, ROI) in each organ/tissue. Appropriate channel colors and names were set for all images (DAPI and DsRed: mCherry). In each annotation area, the total number of cells (DAPI) and the number of DsRed-positive (mCherry-positive) was obtained. The area and average intensity of all the detected cells in each sample was normalized to the background intensity from non-fluorescent cells. The fluorescence intensity was calculated by adapting the corrected total cell fluorescence formula.^53^

### Vector Genome Quantification

Genomic DNA was extracted from liver, kidney, lung, and QUAD muscle using EZNA Tissue DNA kit (Omega Bio-Teck, Norcross, GA, USA) according to the manufacturer’s protocol. Vector genome copies number (VGCN) was measured using a TaqMan qPCR master mix with primers specific for WPRE element sequence of the AAV vector (forward primer: 5’-GGCTGTTGGGCACTGACAA-3’; reverse primer: 5’-CCAAGGAAAGGACGATGATTTC-3’; probe (FAM): 5’-TCCGTGGTGTTGTCG-3’). qPCR assay was normalized using the mouse Titin gene (forward primer: 5’-AAAACGAGCAGTGACGTGAGC-3’; reverse primer: 5’-TTCAGTCATGCT GCTAGCGC-3’; probe (VIC): 5’-TGCACGGAAGCGTCTCGTCTCAGTC-3’).^54^ The VGCN per cell was calculated multiplying WPRE copies normalized on the mouse titin copies for the ploidy of the cells (n = 2).

### *In situ* Hybridization

Tissue cryosections were mounted on SuperFrost Plus Adhesion Slides (Thermo Fisher Scientific, Waltham, MA, USA), with a hydrophobic barrier created using Immedge Hydrophobic Barrier Pen (Vector Laboratories, CA, USA) as previously described^13^. Briefly, RNA staining was performed using the RNAscope multiplex fluorescent reagent kit (Advanced Cell Diagnostics [ACD], Newark, CA, USA) following the manufacturer’s protocol. The AAV RNA was detected with a mCherry probe (ACD, catalog no. 431201-C2). Sequential hybridization procedures were performed with pre-amplifier, amplifier, and label probe. Signal was detected with a tyramine signal amplification (TSA) Plus fluorescein system (PerkinElmer, Waltham, MA, USA). Cell nuclei were stained using manufacturer’s supplied DAPI for 30 s at RT. Finally, slides were mounted with Vectashield mounting medium (Vector Laboratories, Burlingame, CA, USA) and imaged on a confocal microscope using a 20X objective.

### Statistical Analysis

Two-tailed Student’s t test was used to compare between two groups. To compare among multiple groups, one-way ANOVA was used followed by Tukey’s multiple comparisons test. A p value <0.05 was statistically significant. Data are represented as the mean with SEM. Statistical analysis was performed with the GraphPad Prism 9.0 software (GraphPad, La Jolla, CA, USA).

## Author contributions

C.J.M. Investigation, Methodology, Data curation, Formal analysis, Software, Visualization, Writing – original draft; A.C. and J.V. Investigation, Methodology, Data curation, Formal analysis, Writing – review & editing; C.J.M. and E.A.E. Conceptualization, Supervision, Project administration, Funding acquisition, Validation, Writing – review & editing.

## Acknowledgments

We thank Professors Sam Wang for kindly sharing consumables and equipment. This work was partially supported by the PNI Research Innovator grant, Princeton IP Accelerator grant, NJ-ACTS UL1TR003017grant and NIH grants P40OD010996 & NIH-1U01NS113868 (EAE).

